# Multi-Model Machine Learning for Automated Identification of Rice Diseases Using Leaf Image Data

**DOI:** 10.1101/2024.07.09.602645

**Authors:** Rovin Tiwari, Jaideep Patel, Nikhat Raza Khan, Ajay Dadhich, Jay Kumar Jain

## Abstract

**Purpose:** Rice is grown almost everywhere in the world but is notably prevalent in Asian nations where it serves as the main food source for nearly half of the world’s population. Yet, enduring agricultural problems like various rice diseases have been a problem for farmers and planting specialists for ages. A fast, efficient, less expensive, and reliable approach to detecting rice diseases is urgently required in agricultural information since severe rice infections could result in no harvest of grains. Automated disease monitoring of rice plants using leaf images is critical for transitioning from labor-intensive, experience-based decision-making to an automated, data-driven strategy in agricultural production. In the modern era, Artificial Intelligence (AI) is being widely investigated in various areas of the medical and plant sciences to assess and diagnose the types of diseases.

**Methods:** This work proposes a hybrid deep-machine learning system for the automated detection of rice plant diseases using a leaf image dataset. Benchmarked MobileNetV2 architecture is employed to extract the deep features from the input images. Obtained features are fed as input to various machine learning classifiers with different kernel functions using a 10-fold validation strategy.

**Results:** The developed hybrid system attained the highest classification accuracy of 98.6%, specificity of 98.85%, and sensitivity of 97.25% using a medium neural network. The results demonstrate that the established system is computationally faster and more efficient. The proposed system is ready for testing with more databases.

**Conclusions:** The suggested technology accurately diagnoses various rice plant illnesses, reducing manual labor and allowing farmers to receive prompt treatment. Future research topics include incorporating cloud-based monitoring for leaf image capture in non-connected farms, as well as building mobile IoT platforms for continuous screening.

## 1. Introduction

The foundation of the Indian economy is agriculture. In India, farming supports close to 70% of the population [1]. The rural populations of India are reliant on agriculture. A vital source of income for more than 58% of rural residents is farming [2]. Rice cultivation is extremely common in Asian countries, where it is a staple of agricultural and nutritional activities. Rice cultivation takes up a substantial percentage of fertile land in Asia, and millions of farmers work to produce it. The crop’s resilience to a variety of temperatures, from tropical to temperate, helps to explain its extensive cultivation across the continent. Rice is a critical component of food security in many Asian countries, providing millions of people with their primary supply of calories and nutrients. It is the foundation of traditional diets and cultural cuisines, representing nourishment and prosperity [3]. Rice is one of the most important crops in India, but various rice diseases kill roughly 10% to 15% of the rice harvest [3]. Farmers have a long-standing agricultural difficulty in rice cultivation: efficiently managing and reducing illnesses that harm rice plants. Despite the crucial need of good crop care, many farmers lack the necessary expertise and resources to properly identify and manage diseases such as leaf blasts, brown spots, and bacterial blight. These diseases have a severe influence on rice plant growth and productivity, threatening food security and livelihoods. Early detection of infections is critical to minimizing the harm caused by these diseases, but it takes time, skill, and ongoing plant monitoring. Farmers frequently rely on manual observation, but it can be unreliable, time-consuming, and impractical, especially on large-scale farms. As a result, there is a critical need for disease diagnosis and management technologies that are both accessible and efficient to support sustainable rice production practices and strengthen farmers’ resilience to agricultural difficulties [4].

The majority of farmers, however, are less knowledgeable about how to protect plants from different diseases including leaf blasts, brown spots, bacterial blight, etc [4]. These diseases can impact rice plant growth at any stage of production. To avoid major harm, early infection diagnosis is crucial during the early stages of production. It took time, expertise, and ongoing plant monitoring to accurately identify and classify disease in rice plants [5]. Most farmers monitor plant health concerns with their naked eyes, just like a professional would. But, when working with vast farms, manual observation is less reliable, time-consuming, and expensive [6]. To classify and diagnose different diseases at an early stage and with higher accuracy, an autonomous method is needed. To resolve these problems, this study presents a hybrid deep-machine learning system that can determine the state of the plant’s health using image artificial intelligence tools. Every disease has symptoms that appear on the leaf, stems, or fruits. With the aid of deep-machine learning methods, the observation of leaf images can be used to detect diseases in rice plants by scanning for these symptoms. The algorithm recognizes the condition of the rice plant when there are unhealthy leaves. Our suggested model focuses on the bacterial blight, brown spot, and leaf blast of rice. This study includes three most common rice diseases namely, brown spot, bacterial blight, and leaf blast. Brown spot disease is a fungal infection that commonly damages rice plants. Bipolaris oryzae or Helminthosporium oryzae is the fungus that causes it. The disease causes small, oval to elliptical spots to form on the leaves, leaf sheaths, and panicles of rice plants. Initially light brown, the spots darken and have a yellowish halo as the condition advances. The affected leaves may break off, curl, and dry out [7]. Another deadly rice plant disease is called bacterial blight, which is brought on by the bacterium Xanthomonas. When water-soaked lesions emerge on the leaves, stems, and grains of rice plants, it’s time to identify the illness. Lesions gradually turn brown and yellow as the sickness progresses, causing the damaged tissue to die [8]. The rice-damaging fungus Pyricularia oryzae is the source of the fungal disease known as leaf blast. It is one of the worst diseases that affect rice, resulting in large losses in output. The illness causes tiny, round-to-elliptical lesions on the leaves, which may have a pale green or yellow halo surrounding them [9]. It is typical to see lesions with a reddish-brown ring surrounding a grey-to-whitish center. As the condition worsens, the lesions may combine to form large, asymmetrical lesions that can seriously damage the leaves. These illnesses are frequently more severe in areas with high humidity, frequent rains, and warm temperatures. If left untreated, the illness can spread swiftly and cause significant losses in yield. The application of cultural control techniques, such as planting disease-resistant cultivars and upholding proper field hygiene, can reduce the disease’s incidence and severity. These diseases can also be managed chemically by using antibiotics and copper-based fungicides, although their efficacy may be compromised by the growth of bacterial strains that are resistant to these treatments. Early detection of these diseases is necessary to protect the crops from the viruses. To successfully cultivate crops, it is essential to accurately identify the symptoms of plant leaves, identify the related pest or disease, and calculate the proportion of the disease. Only a small number of urban agricultural facilities manually identify plant illnesses at this time. It is common for farmers in rural areas to have to travel great distances to discover crop diseases. However, our system is an automated approach that uses input photos of sick rice plants to diagnose diseases in rice plants. The system produces an extensive report that includes symptoms and suggestions for managing the detected illness. In the literature, researchers have proposed various techniques to detect these infections using leaf images.

Singh et al. [10] used image processing techniques to identify plant diseases. Many segmentation and classification methods were used with MATLAB to forecast diseases in a range of crops, such as potatoes, tomatoes, jackfruit, bananas, mangoes, beans, and lemons. A genetic algorithm was used in the segmentation procedure. Additional techniques like Artificial Neural Network (ANN), Fuzzy Logic, Bayes classifier, and hybrid algorithms were investigated to improve the recognition rate during classification. A thorough investigation of plant disease identification using image processing was provided in Khirade and Patil’s work [11]. A range of feature extraction strategies, segmentation tactics, and classification algorithms were covered throughout the conversation. Boundary and spot recognition, Otsu threshold, and k-mean clustering techniques were among the segmentation algorithms used. Leaf color evaluation and co-occurrence approach by utilizing H and B components were included in the feature extraction algorithm. Likewise, ANN, Backpropagation Neural Network (BPNN), modified Self-Organizing Map (SOM), and Multiclass SVM were among the classification techniques investigated in the study. By applying image processing techniques, the integration of these methodologies facilitates the precise diagnosis and classification of various plant diseases. Mahalakshmi and Srinivas used the Firebird V robot to do Vision-based Plant Leaf Disease Detection using Color Segmentation in their work [12]. For the investigation, a high-quality camera-equipped Firebird V Robot was used to take pictures of different plants. Neural networks were then set up in combination with image processing methods to accurately identify the impacted leaf portions. The study shows that there is a need for improvement in the creation of a creative, effective, and quick interpretation system for diagnosing plant diseases. The author used image processing techniques in a systematic review to detect agricultural diseases in [13]. The study outlines several methods for picture pre-processing, image segmentation, feature extraction, and image classification. Two popular image pre-processing methods are contrast enhancement and noise filtration. Similarly, K-means, region expanding, foreground-background, threshold-method, morphological operators, fuzzy c-mean, color segmentation, and canny edge detection are among the most common image segmentation techniques mentioned in the review. The study also examines various classification methods, including Radial Basis Function (RBF), K-Nearest Neighbors (KNN), Artificial Neural Network (ANN), Support Vector Machine (SVM), and Backpropagation Neural Network (BPNN). The author suggests that researchers can leverage these methods to achieve a high level of accuracy in their work. An overview of several classification and segmentation methods for the identification of plant diseases was given by Khan and Oberoi in [14]. Partition clustering, edge detection, and region-based segmentation are the three main segmentation strategies that the authors discovered. Subtypes of region-based segmentation included region growth, region splitting, and region merging. The study also covered neural network-based classification techniques, such as Singular Value Decomposition (SVD), Principal Component Analysis (PCA), and backpropagation analysis. A review has been conducted by Shruthi, Nagaveni, and Raghavendra [15] on the various methods of machine learning for plant disease diagnosis. Researchers outlined and contrasted several fundamental classification methods for identifying the signs of plant diseases. A. K. Rath and J. K. Meher emphasized the effect of plant diseases on agriculture output and the country’s economy in [16]. Acknowledging the need for prompt and precise identification, the authors utilized image processing methods to distinguish and categorize two rice illnesses, specifically rice blast and brown spot. For detection and classification, the implemented approach made use of a classifier known as a Radial Basis Function Neural Network (RBFNN). Strong generalization, online learning, and a straightforward design are among the advantages of the RBFNN [17]. In the domain of disease prediction challenges, In [18] confirmed RBFNN is very efficient and dependable when compared to other networks. The database, which included pictures of rice blast illness, rice spot disease, and healthy leaves, was created and kept up to date by the writers. There were about 100 photos in each lesson, for testing and instruction. The authors of [19] presented a unique method for detecting illnesses in rice plants by combining Faster R-CNN and FCM-KM techniques. The study tackled a range of obstacles related to illness identification in rice plants, utilizing distinct methods and filters to alleviate these issues. A weighted multi-level median filter with a 2-dimensional mask has been used to reduce image data noise. Otso threshold segmentation algorithm was employed by the authors to address complicated background interference. To classify diseases, the R-CNN algorithm was utilized. The authors claimed that their method for identifying these illnesses is not only quicker but also more capable. They also emphasized how their approach might be used with mobile devices and the Internet of Things (IoT) to identify plant illnesses in real-time. Mobile IoT platforms play an important role in the screening of rice plant diseases by enabling real-time monitoring and data collection. These platforms combine multiple sensors, like as cameras and environmental sensors, to collect detailed information about plant health and environmental conditions. A deep CNN model was used by Wan-Jie Liang et al. in [20] to identify rice blast illness. In their investigation, the scientists used a dataset that included rice blast photos that were both infected and uninfected. Jiangsu Academy of Agricultural Sciences China of Plant Protection provided them with 2902 pictures of healthy rice plants and 2906 images of infected rice leaf samples, which depict the rice blast problem. The author provided evidence that the CNN method works better than others and can be used in real-world situations. CNN efficiently extracts blast disease-related features and then categorizes the symptoms of the illness. The creation of the rice blast disease dataset is one of this study’s major contributions. This dataset was integrated with other publicly available datasets by the author to create an extensive dataset. Afterward, this dataset was used by other researchers to investigate the identification of rice plant diseases.

In this work, an innovative approach for automated detection of rice plant diseases is proposed. Initially, deep features are extracted from leaf images using pre-trained models, specifically MobileNetV2, DarkNet19, and ResNet18 convolutional neural networks (CNNs). These models are chosen for their effectiveness in image feature extraction. Subsequently, the extracted features from the three CNNs are merged to create a comprehensive representation of the input images. To classify the images and identify infected plants, machine learning classifiers are employed. Three classifiers are utilized for this task: ensemble classifiers with various kernel functions, k-nearest Neighbors (KNN), and Support Vector Machine (SVM). The proposed system can contribute to reducing manual labor in disease diagnosis by automating the detection of rice diseases through the classification of leaf images. This automation streamlines the diagnosis process, potentially minimizing the need for manual inspection and analysis by agricultural experts. The key contributions of this study are as follows:

1. We propose a new hybrid deep-machine learning approach that is intended to quickly and precisely identify rice illnesses. Using the best features of both paradigms, this novel method combines CNN with conventional machine-learning techniques to improve the accuracy and efficiency of rice plant disease identification.
2. The method uses a CNN to automatically identify subtle patterns linked to different diseases by extracting complex information from photos of rice plants. To complete the final classification, these deep features are then input into machine learning algorithms like ensemble classifiers, KNN, and SVM. The process of identifying diseases is made comprehensive and reliable by the synergistic blending of deep learning and conventional machine learning approaches.
3. Our hybrid approach seeks to enable fast and accurate identification of rice diseases by combining the interpretability and generalization qualities of classical machine learning methods with the image representation capabilities of CNNs. This method takes advantage of the well-established machine learning concepts for efficient illness classification decision-making, in addition to harnessing the power of deep learning for feature extraction.
4. Produce improved classification performance that outperforms the traditional methods.

The remaining section of this paper is organized as follows: Section 2 includes the materials and methods; Section 3 offers the results of the simulation; Section 4 shows how this work is discussed with previous studies; and Section 5 concludes and shows where this study will go in the future.

## 2. Materials and Methods

The presented study describes the outcomes of a real-world implementation of an artificial intelligence (AI) system for detecting rice diseases using leaf images. A publicly available dataset with a 10-FCV scheme is used to validate the presented approach. The transfer learning method is employed to re-train the utilized deep learning models. Deep features are extracted using three benchmark CNNs. Deep features of all three CNNs are merged and classified using three machine learning algorithms with different kernel functions. Figure 1 depicts the schematic view of the presented system.

**Figure 1:**
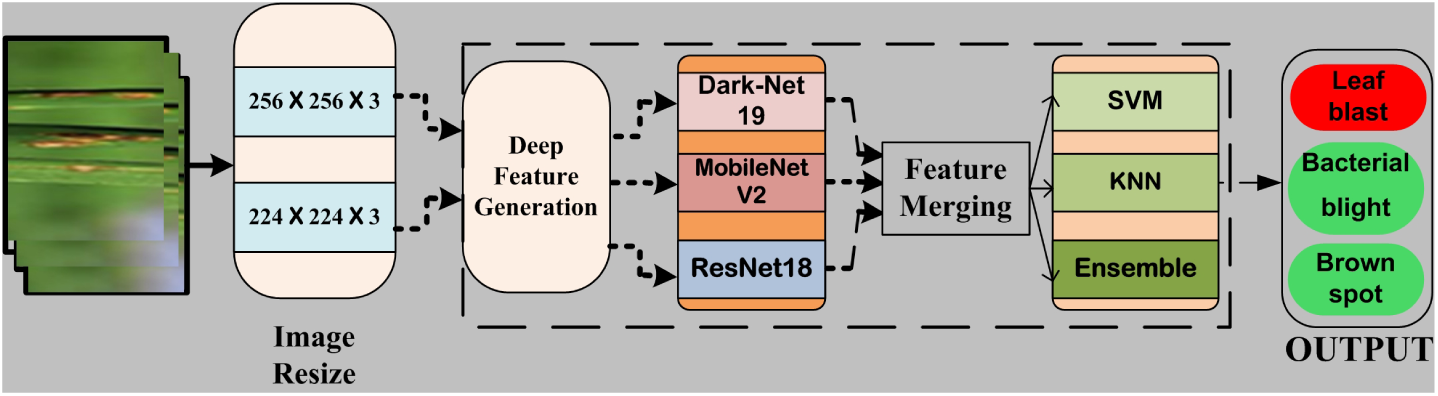
Schematic view of the presented system.

### 2.1. Dataset

The leaf images utilized in our research were gathered from various reputable sources, including the UCI and Kaggle datasets, along with specific images obtained from the IEEE dataset repository, which encompassed both disease and plant categories. To ensure the compatibility and harmonization of these disparate sources, several steps were undertaken. Firstly, we conducted meticulous preprocessing and standardization procedures to homogenize the images in terms of resolution, format, and quality. Additionally, rigorous quality control measures were implemented to filter out any inconsistencies or anomalies within the dataset. Furthermore, during the dataset compilation phase, particular attention was paid to maintaining a balanced representation of the three distinct disease classes: bacterial blight, leaf blast, and brown spot. This meticulous curation process aimed to mitigate any potential biases or disparities that could arise from the amalgamation of heterogeneous sources. We included, in particular, 1030 images of rice spot, 1130 images of rice blast, and 1536 images of rice blight disease. Six randomly chosen photos of rice leaves, each representing a different illness, are shown in Figure 2.

**Figure 2:**
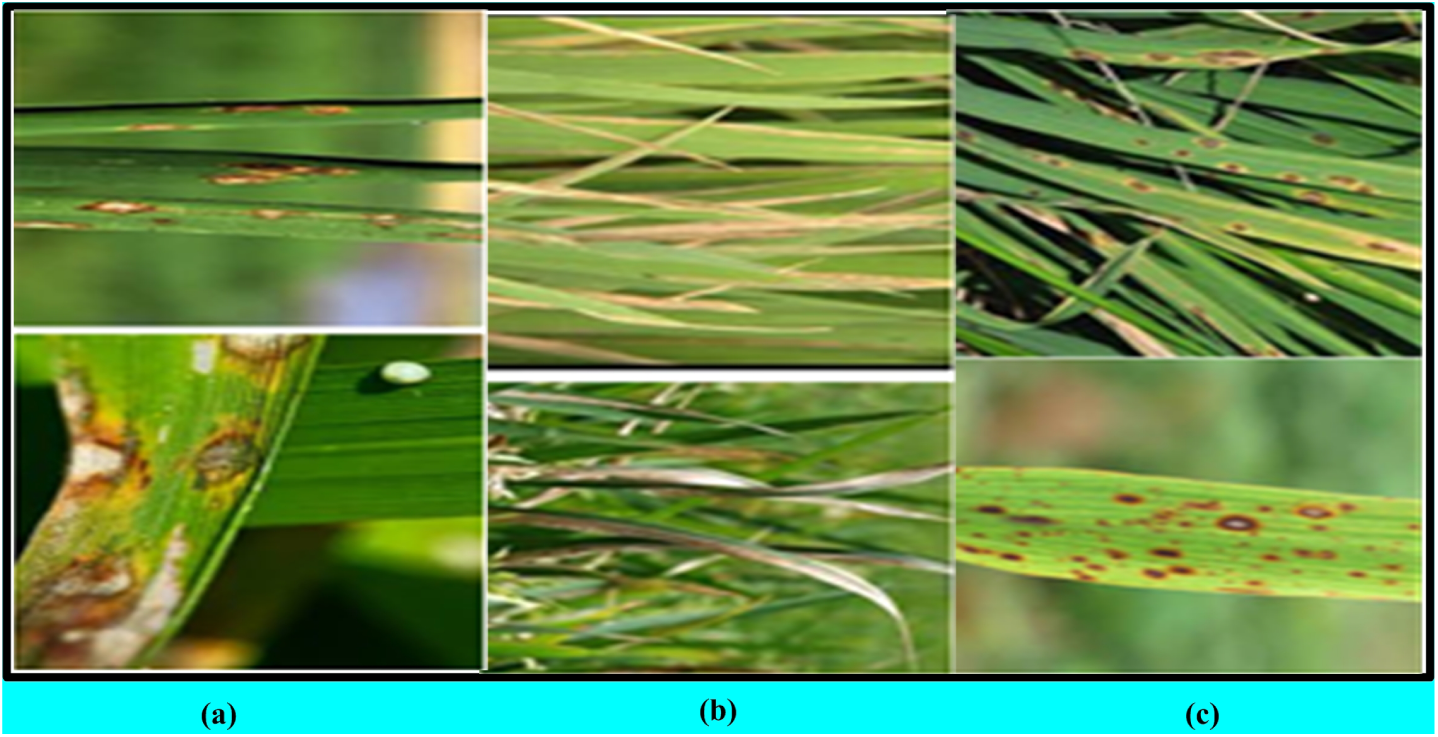
Rice-leaf images of (a) Brown Spot Leaf, (b) Bacterial Blight, and (c) Blast Disease

#### 2.1.1. Verify the Image Clarity

After images are collected, their quality is evaluated. When the input image quality satisfies the predetermined standards, noise reduction preprocessing is started, and the input matrix of the model is put through quality assessment. When an image is rejected by the model and a corresponding message is displayed, it means that the image quality is not up to par. To assess the quality of input images, CNN uses the BRISQUE algorithm that is built into the ’Imagequality’ package. CNN libraries use the scores that the BRISQUE algorithm produces for each input image to evaluate the quality of the image. Through transformative feature computations, this approach, which uses image pixels to compute features, is highly effective in assessing the quality of photographs of rice plants.

### 2.2. Convolutional Neural Network (CNN)

CNN-based frameworks are widely studied in current research, especially for various classification challenges. CNN’s attraction is its capacity to substitute an automated system for conventional, laborious approaches including feature evaluation, selection, and classification. The process is streamlined and made more flexible by this automated method. CNN is capable of performing a wide range of pattern recognition tasks, including object localization, segmentation, detection, and image classification. Because of its adaptability, CNN-based frameworks have been widely used and investigated in a variety of disciplines [21]. These networks are built using a technique called Representation-Based Learning (R-BL). The R-BL system is a systematic collection of procedures meant to allow a CNN to automatically obtain appropriate representations for classification after receiving relevant input. The model in the R-BL framework selects pertinent features on its own and combines the feature selection and classification phases into a single pipeline with ease. This simplified method improves the network’s performance by enabling automatic learning of representations that are suitable for classification tasks. Thus, the R-BL technique helps to create CNNs in a more efficient and well-organized manner, where the system takes care of the intelligent feature extraction, selection, and classification [22]. Deep learning approaches are used to diagnose rice leaf illnesses. These approaches include deep feature extraction, transfer learning to fine-tune benchmark Convolutional Neural Networks (CNNs), and extensive training and validation of the proposed framework. All of these techniques work together to make the model more successful in precisely identifying and categorizing different rice plant illnesses. Furthermore, the use of deep learning has demonstrated its effectiveness in a variety of biomedical tasks, such as segmentation issues, diagnosis of various diseases, and signal and image categorization. The adaptability shown in these applications highlights the potential of deep learning techniques, which makes them useful instruments for dealing with difficult problems in a variety of fields, such as plant pathology and agriculture [23]. One type of representation-based learning (RBL) that has a structured design is called deep learning. An input layer, several convolutional layers (Conv2DL), pooling layers (PL), a few fully connected layers (FCL), and an output classification layer are the usual components of this architecture. Together, the essential elements of this architecture allow the model to automatically extract hierarchical representations from the input data; this makes deep learning very useful for intricate tasks like image categorization and recognition. Convolution operations are carried out by the Conv2DL layers to capture spatial patterns; dimensionality is reduced by the pooling layers; and the fully connected layers allow the model to establish complex correlations between features for precise classification in the output layer. Deep learning stands out as a potent paradigm in many domains, such as pattern recognition and computer vision, thanks to its hierarchical and automated representation learning [23]. In this research, three benchmarks CNNs, DarkNet19, ResNet18, and MobileNetV2, are used for disease detection using leaf images.

A sophisticated CNN model created especially for object detection is called Darknet-19. It is a crucial part of the object detection models created by Joseph Redmon and Ali Farhadi as part of their YOLO (You Only Look Once) series. When it comes to real-time object detection tasks, the YOLO models are renowned for their effectiveness and quickness [24, 25]. With 19 convolutional layers, Darknet-19 is a lighter variant of the YOLO series models, which makes it appropriate for situations when resources are limited. Darknet-19 retains a high degree of accuracy in object detection tasks even with its decreased complexity.

The architecture of the model utilizes several convolutional layers to extract features, which facilitates the analysis and identification of objects in a picture. With models like Darknet-19, the YOLO framework has been widely applied in a variety of fields, such as robotics, autonomous vehicles, and surveillance, where precise and real-time object detection is essential. Google has created a small CNN model called MobileNetV2, which is intended for effective and efficient object recognition and classification in embedded and mobile applications. It is the replacement for the initial MobileNet model, which was created to offer a thin CNN model that could function well on portable devices. Because MobileNetV2 is built on an inverted residual structure, it can achieve good accuracy at low computational complexity and small model sizes. It can carry out image classification and object recognition tasks because it was trained on the ImageNet dataset, which is a sizable collection of images annotated with various item types. MobileNetV2’s effective use of computing resources is one of its primary characteristics, which makes it a good choice for deployment on mobile and embedded devices with constrained memory and processing power. It is ideal for applications like video surveillance and autonomous cars since it can operate in real-time on these devices. An input image with dimensions of 224 x 224 is used by MobileNetV2. There are two more features in MobileNetV2. The first kind of bottlenecks are linear ones. The quickest route between bottlenecks is the second. MobileNetV2 can be used for object detection, feature generation, and classification. A vast collection of images annotated with various item types makes up the ImageNet dataset, which is used to train the 18-layer CNN known as ResNet18. By using an image as input and producing a set of class probabilities for each object it identifies in the image, it may carry out tasks related to object identification and image classification. Microsoft Research created the ResNet18 model, which is intended for use in object identification and picture classification applications. This model is a variation of ResNet, which was created to help with the difficulty of training extremely deep CNNs. The 224 x 224 input image is accepted by the network.

### 2.3. Features extraction

In classical machine learning, feature extraction is an essential stage that involves finding and extracting pertinent information from unprocessed data to improve the model’s capacity for classification and prediction. According to the mentioned work [26, 27], Convolutional Neural Networks (CNNs) are especially good at feature extraction because they can automatically train and extract hierarchical features over numerous layers, enabling robust classification capabilities. However, when working with big datasets, training CNNs from scratch can be computationally taxing and time-consuming. Transfer learning is used to address this problem. Transfer learning lessens the computational load of feature extraction by utilizing training data from one task and applying it to another that is similar. Three benchmark deep models are used as pre-trained feature generators in this work: ResNet18, MobileNetV2, and DarkNet19. These models can capture rich and generalizable properties since they have been trained on large datasets for a variety of tasks. The time and resources needed for training from scratch are greatly decreased by employing these pre-trained models. ResNet18 is a variation of the ResNet architecture that is well-known for its efficacy in image classification tasks; MobileNetV2 is specifically intended for efficient processing on mobile and embedded devices; and DarkNet19 is a convolutional neural network designed for object detection. The study intends to leverage the learned representations from other domains to improve the model’s capacity to categorize and identify patterns in the provided data by integrating these pre-trained models for feature extraction.

### 2.4. Classification of features using traditional machine learning Methods

The potential of learned image features for rice disease identification is being researched in addition to the application of the transfer learned network for screening [28]. The network activations from the various learned model layers are collected and fed into the SVM, KNN, and Ensemble-proven state-of-the-art classifiers. In this paper, a 10-FCV approach is used to construct the model, which helps to minimize potential bias during model generation. The efficacy of the suggested feature extraction model is assessed using the classification results obtained. Brief details for SVM, KNN, and Ensemble classifiers are summarized as follows: SVM is a form of supervised machine learning method that can be used for classification, regression, and outlier detection. They work by identifying a hyperplane in a high-dimensional space that isolates dissimilar classes as much as possible [29]. SVMs excel at managing high-dimensional data and complex decision boundaries in the context of classification, displaying robustness to overfitting and making them acceptable for jobs with small or moderate-sized datasets. To improve SVM’s flexibility in capturing varied patterns, this work employs five kernel functions: linear, cubic, quadratic, coarse Gaussian, and medium Gaussian. K-Nearest Neighbors (KNN) is a simple and effective method for classification and regression tasks that uses the instance-based learning principle. It saves training data and predicts future instances based on their similarity to previously saved data [30, 31]. KNN, which is frequently used for nonparametric classification, selects the ideal *k* (number of neighbors) value and assigns labels based on the most common labels from the *k*-nearest neighbors. While KNN is simple and flexible, its computational complexity can be a disadvantage for large datasets. To increase the versatility of the KNN approach, five distinct kernel functions are used in this study: Cosine, Cubic, Fine, Weighted, and Medium. When compared to a single estimator, ensemble approaches integrate predictions from numerous base estimators to improve generalizability and robustness [32]. The goal is to improve model generalization by decreasing overfitting and increasing prediction stability. This research employs four ensemble methods, including Bagging, Subspace, RUSBoosted, and Subspace, to leverage the collective strength of various base models for superior task performance.

### 2.5. Performance Metrics

The developed hybrid system achieves several key performance metrics to assess its effectiveness. These metrics include accuracy (*A_CC_*), sensitivity (*S_EN_*), specificity (*S_P E_*), F-1 measure, precision (*PRC*), negative prediction value (*N_P V_*), and area under the curve (*AUC*). Each of these metrics provides valuable insights into different aspects of the system’s performance, such as its ability to correctly classify both positive and negative instances, its precision in identifying true positives, and its overall discriminatory power. By considering multiple performance characteristics, the proposed approach aims to comprehensively evaluate its superiority and effectiveness in automated rice plant disease detection. The mathematical representation of these performance characteristics is given as [33, 34]:

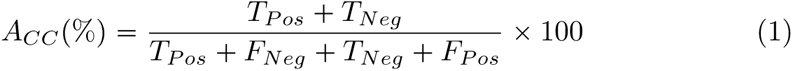

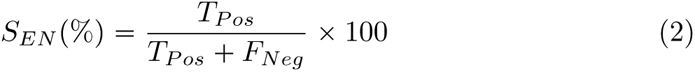

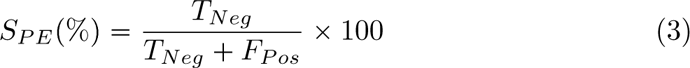

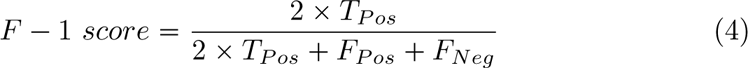

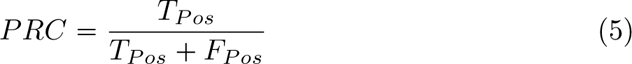

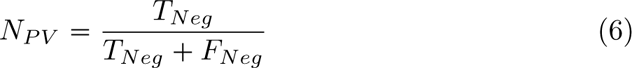

where, true positive (*T_P os_*), false positive (*F_P os_*), true negative (*T_Neg_*), and false negative (*F_Neg_*) are the confusion matrix parameters.

## 3. Simulation Setup and findings

The experiment is run on a single CPU with a Core i7 processor, 24-GB RAM, a 1-TB SSD, and a 512-GB hard disk, using the MATLAB platform. To augment the dataset and enhance its size, we employed a combination of common data augmentation techniques. These included rotation, reflection, and shearing of the input images (randomly rotated from -5^0^ to 5^0^, a reflection of 1, and a range of horizontal-vertical shear is ([-0.05, 0.05]). Rotation involved rotating the images by various angles, while reflection encompassed flipping the images horizontally and vertically. Additionally, shearing was applied to introduce geometric transformations by distorting the images along their axes. By systematically applying these augmentation methods, we effectively diversified the dataset, enriching it with variations of the original images to enhance model robustness and generalization capabilities. A 10-fold cross-validation (10-FCV) scheme is used for a more robust evaluation. This approach involves dividing the extracted features into ten equal parts, with one part reserved for testing, one for validation, and the remaining eight for model training. The process is repeated ten times, each time using a different partition for testing and validation while utilizing the remaining partitions for training. This strategy ensures comprehensive evaluation and validation of the model’s performance across multiple iterations, enhancing its reliability and generalization capability. The Adam optimizer is used with a starting learning rate of 0.0001 to maximize the learning rate and reduce cross-entropy loss. Table 1 shows the hyperparameter settings for the second and third studies. This comprehensive setup ensures that the proposed framework is thoroughly evaluated, taking into account both hardware specifications and robust cross-validation processes. Figure 3 shows the training progress graphs yielded for (A) MobileNetV2 (B) Resnet-18 (C) DarkNet19. Table 2 gives the overall classification accuracy yielded by classifying features extracted by individual CNNs and merged features of all CNNs, using various machine learning algorithms. It is evident from Table 2 highest classification accuracy of 98.9% is achieved by classifying merged features of all CNNs by applying the SVM algorithm with a medium Gaussian kernel function. Various performance metrics are evaluated, to check the effectiveness of the presented SVM algorithm with medium Gaussian kernel function to classify merged features of all CNNs into different disease classes. Figure 4 shows the bar chart of the obtained classification accuracy using different kernels of the machine learning classifier. Table 3 gives the overall performance metrics obtained for the SVM algorithm with a medium Gaussian kernel to classify merged features. It can be noted from Table 3, that the SVM classifier with a medium Gaussian kernel has achieved the *S_EN_* of 97.25%, *S_P E_* of 98.85%, *F* 1-score of 97.2, *PRC* of 98.6, and *N_P V_* of 97.8 in classifying merged features of all three explored CNNs. Figure 5 shows the pie chart of the obtained classification performance.

**Figure 3:**
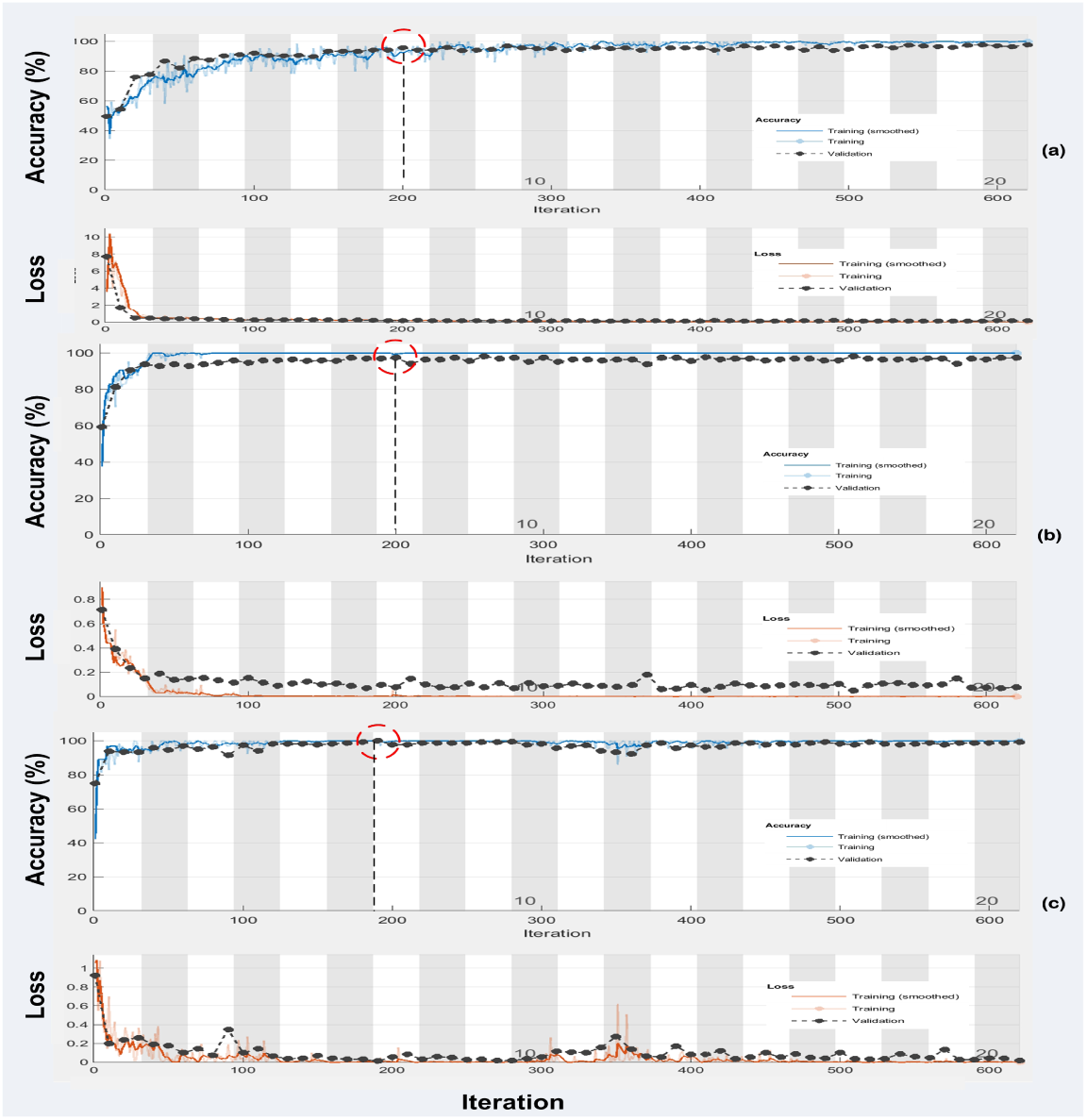
Training progress graphs yielded for: (A) MobileNetV2 (B) Resnet-18 (C) DarkNet19

**Figure 4:**
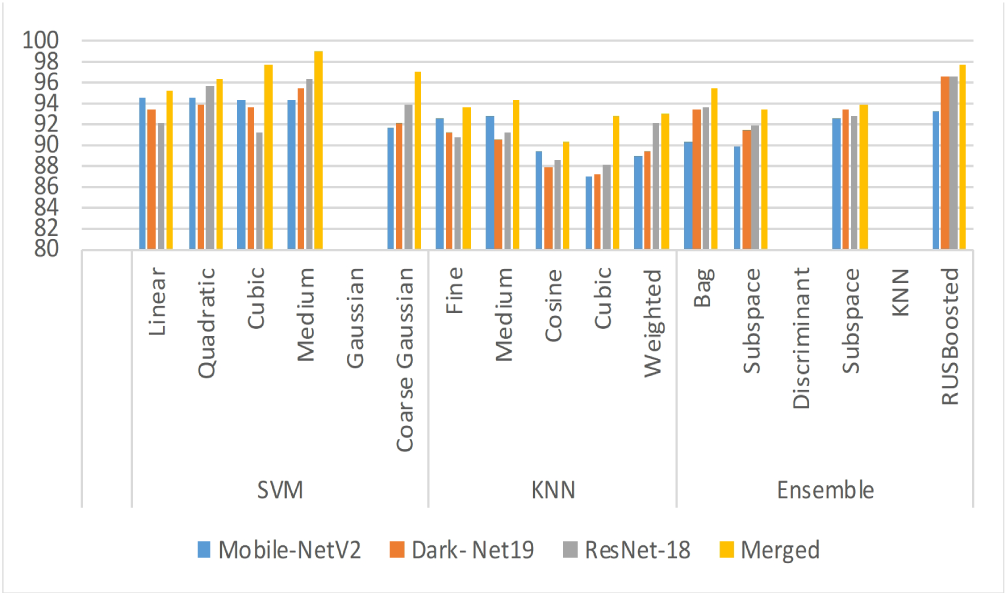
Bar chart of the obtained classification accuracy

**Figure 5:**
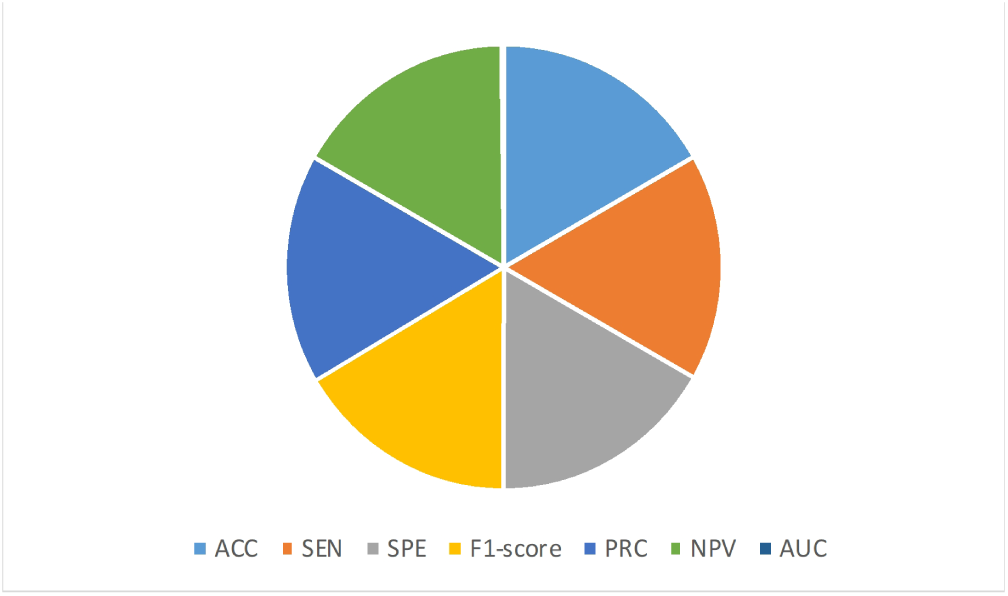
Pie chart of the obtained classification performance SVM classifier with medium Gaussian kernel.

**Table 1:** Details of hyperparameters used in second and third experiments.

| **Classifier** | **Kernel**<br>**Function** | **Box**<br>**Constraint**<br>**Level** | **Kernel**<br>**Scale** | **Multi-**<br>**class**<br>**Method** | **Stand-**<br>**ardize**<br>**Data** |
| --- | --- | --- | --- | --- | --- |
| SVM | Linear<br>Cubic<br>Quadratic<br>Coarse<br>Gaussian<br>Medium<br>Gaussian | 1 | Automatic | One-vs-One | True |
| **Classifier** | **Kernel**<br>**Function** | **Number**<br>**of**<br>**Neighbors** | **Distance**<br>**Metric** | **Distance**<br>**Weight** | **Stand-**<br>**ardize**<br>**Data** |
| KNN | Cosine<br>Cubic<br>Fine<br>Weighted<br>Medium | 10<br>10<br>1<br>10<br>10 | Cosine<br>Minkowski<br>Euclidean<br>Euclidean<br>Euclidean | Equal<br>Equal<br>Equal<br>Squared<br>Inverse<br>Equal | True |
| **Classifier** | **Ensemble**<br>**Method** | **Number**<br>**of**<br>**Split** | **Number**<br>**of**<br>**Learner** | **Learning**<br>**Rate** | **Subspace**<br>**Dimension** |
| Ensemble | Bag<br>Subspace<br>Discriminant<br>RUSBoosted<br>Subspace<br>KNN | 20 | 30 | 0.1 | 1 |

**Table 2:** Summary of classification accuracies (%) obtained using classifiers.

| Classifier | Kernel<br>Function | Mobile-NetV2<br>Features | Dark-Net19<br>Features | Res-Net18<br>Features | Merged<br>Features |
| --- | --- | --- | --- | --- | --- |
| $(A_{CC}\%)$ | | | | | |
| SVM | Linear | 94.6 | 93.5 | 92.1 | 95.2 |
|  | Quadratic | 94.6 | 93.8 | 95.7 | 96.2 |
|  | Cubic | 94.2 | 93.7 | 91.1 | 97.6 |
|  | Medium<br>Gaussian | 94.3 | 95.5 | 96.3 | **98.9** |
|  | Coarse<br>Gaussian | 91.6 | 92.1 | 93.8 | 96.9 |
| KNN | Fine | 92.6 | 91.1 | 90.7 | 93.6 |
|  | Medium | 92.8 | 90.6 | 91.2 | 94.4 |
|  | Cosine | 89.4 | 87.8 | 88.5 | 90.2 |
|  | Cubic | 86.9 | 87.2 | 88.1 | 92.8 |
|  | Weighted | 88.9 | 89.5 | 92.1 | 93.0 |
| Ensemble | Bag | 90.3 | 93.3 | 93.6 | 95.5 |
|  | Subspace<br>Discriminant | 89.8 | 91.4 | 91.9 | 93.3 |
|  | Subspace<br>KNN | 92.5 | 93.4 | 92.7 | 93.9 |
|  | RUSBoosted | 93.2 | 96.5 | 96.5 | 97.7 |

**Table 3:**
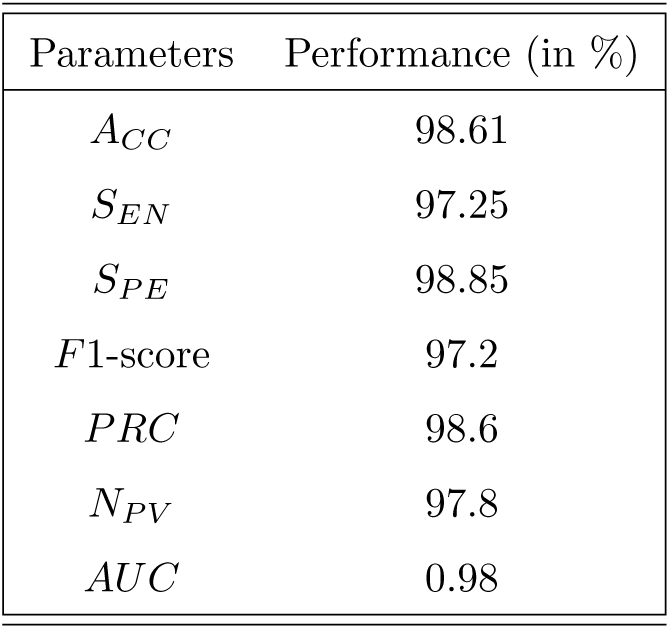
Performance metrics obtained for SVM classifier with medium Gaussian kernel.

| Parameters | Performance (in %) |
| --- | --- |
| $A_{CC}$ | 98.61 |
| $S_{EN}$ | 97.25 |
| $S_{PE}$ | 98.85 |
| $F1$-score | 97.2 |
| $PRC$ | 98.6 |
| $N_{PV}$ | 97.8 |
| $AUC$ | 0.98 |

### SVM classifier with a medium Gaussian kernel

Figure 6 represents the overall receiver operating characteristic (ROC) plots for medium Gaussian SVM. Figure 5 specifies that the classifier obtains a value of 0.98 for True Positive Rate (TPR) vs. false Positive fraction (FPR).

**Figure 6:**
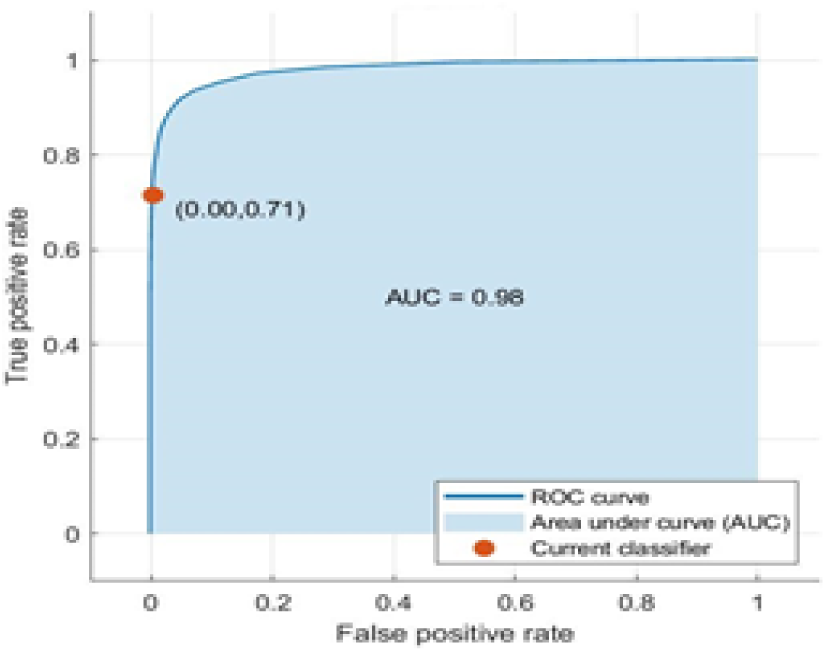
ROC plots achieved for SVM classifier with medium Gaussian kernel.

## 4. Discussion

Plant disease diagnosis and classification from leaf images has emerged as a fascinating area of disease detection at the nexus of computer science and agriculture. In agriculture, a variety of computer vision/artificial intelligence-based methods are utilized to identify and categorize illnesses from images of plants. Farmers encounter difficulties in recognizing infections due to their limited awareness of disease symptoms, which can lead to crop losses. Manual detection is a laborious and imprecise method in which farmers observe plants at every stage of growth. To solve these issues in the context of rice plants, this study makes use of image processing and computer vision, particularly deep hybrid approaches. Unlike existing approaches, our method capitalizes on the complementary strengths of CNNs for image feature extraction and conventional machine learning algorithms for robust classification. By leveraging CNNs to automatically discern intricate patterns associated with various diseases from rice leaf images, we pave the way for more nuanced and accurate classification outcomes. Subsequently, these extracted deep features are seamlessly integrated into ensemble classifiers, KNN, and SVM algorithms, enabling comprehensive disease detection with enhanced reliability. The main objective of this research is to use hybrid deep machine learning to detect three major rice plant illnesses to maximize rice output in terms of both quality and quantity. The main goal is to help rice farmers, analysts, and other stakeholders by offering a precise and timely method for identifying rice plant illnesses. The two stages of the suggested model are feature extraction and classification. During the feature extraction phase, a predetermined collection of pictures of rice plant leaves—including those with bacterial leaf blight, brown spot disease, and leaf blast disease—is used to extract deep features from pre-trained deep learning models. The diagnosis, classification, and cure recommendation processes are under the purview of the disease prediction phase using machine learning models. Based on information obtained from agricultural specialists, the algorithm classifies the type of disease and proposes a specific therapy in addition to predicting whether a leaf is healthy or sick. The proposed hybrid deep-learning model was specifically designed to address the three primary goals outlined in the research:

1. Determine whether a rice leaf is healthy or not: The model successfully achieved this goal by accurately classifying rice leaf images as either healthy or diseased, thereby enabling farmers to identify potential health issues affecting their crops.
2. Identify the sort of disease: Through comprehensive analysis and classification, the model accurately identified various types of rice diseases, including bacterial blight, leaf blasts, and brown spots, among others. This achievement fulfills the second goal of the research, providing farmers with crucial information about the specific diseases affecting their crops.
3. Make a recommendation for how to treat the illness that is anticipated: While the manuscript may not explicitly detail treatment recommendations, the successful identification and classification of rice diseases empower farmers to take timely and informed action to address the identified illnesses.

The proposed technology offers several potential benefits for farmers, particularly in the context of rice plant disease management:

1. The proposed system provides a reliable and efficient method for diagnosing and classifying plant diseases from leaf images, leveraging advancements in computer vision and artificial intelligence.
2. By automating the disease detection process, farmers can overcome challenges associated with limited awareness of disease symptoms, thereby reducing the risk of crop losses and improving overall yield.
3. This study offers a promising solution to enhance disease surveillance and management practices in rice cultivation, ultimately empowering farmers with tools to make informed decisions and optimize crop health and productivity.

## 5. Conclusion and Future Directions

Early diagnosis and accurate treatment of illnesses can greatly reduce crop productivity losses. With the use of the proposed hybrid model for disease prediction, rice crop productivity has increased significantly overall. The model’s predictions direct the use of treatments tailored to a particular condition, including sprays or medications. While many contemporary image processing techniques help phytopathologists and farmers anticipate plant diseases early on, these techniques frequently fall short of providing a thorough examination of rice plant diseases. Furthermore, they might not be able to diagnose patients promptly, which would restrict farmers’ ability to prevent illness and lessen its possible effects. To address the weaknesses in the current methods, we thoroughly reviewed the literature, found flaws, and learned the fundamentals of illnesses that impact rice harvests. Our suggested method makes use of CNN and machine learning modules to identify impacted areas of leaves on rice plants. The technology counts the pixels in the affected area to ascertain the proportion of the leaf afflicted after this detection and analyzes the disease to establish its type. The main conclusions show that our approach makes it simple and inexpensive for farmers to identify the three main rice plant illnesses. Our method provides farmers and agriculturists with a useful tool by automating the manual disease detection system using proposed techniques, allowing for the quick prediction of the three primary illnesses that affect rice plants. This increases crop productivity while also saving time and resources. Furthermore, this approach is adaptable enough to add to or enhance other functions. The paradigm can be adjusted to suit changing standards and problems as user demands for increasingly efficient and user-friendly interfaces develop in line with emerging technology. Our research objectives have been successfully met, and the classification model we created is an example of a conceptual framework that has been applied in a variety of ways to meet the needs of farmers. While the suggested methodology has demonstrated impressive results, there are several avenues for future exploration. The proposed method could be extended to include disease prediction in various other crops beyond the scope of this study. This expansion would contribute to a broader understanding of agricultural challenges and the applicability of our approach across different agricultural contexts.

